# Ambient temperature storage of individual parasitic nematode larvae for whole-genome sequencing

**DOI:** 10.64898/2026.05.15.724465

**Authors:** The WORMS-SEA Consortium, Erwan Budi Hartadi, Elsa Herdiana, Hesham Mahyoub Al-Mekhlafi, Arutchelvan Rajamankam, June Haidee Acūna-Lariosa, Jennifer Luchavez, Vanessa Joy Mapalo, Modupeh Betts, Stephen Doyle, Mark Viney

## Abstract

Soil-transmitted helminth (STH) infections are a major public health burden, and there are programmes of mass drug administration that attempt to ameliorate the harm that they cause. There has been increasing use of genomics to study STH infections and other parasitic nematodes, with particular interest in whole genome sequencing (WGS). For such studies, samples are commonly stored frozen, but in settings where these infections are endemic this can be difficult, and so there would be advantages to having ambient temperature storage methods. We investigated two ambient temperature storage methods – FTA cards and DESS buffer – for infective larvae of the rat parasites *Nippostrongylus brasiliensis* and *Strongyloides ratti*, prior to DNA extraction and then WGS. Our results showed that for individual larvae stored on FTA cards or in DESS buffer, this resulted in a lower proportion of sequence reads that mapped to the reference genomes, compared to the frozen control samples. Generally, for individual larvae, DESS-storage resulted in better sequencing results than FTA-storage. However, for pools of 10 or 50 larvae, then these ambient temperature storage methods generally resulted in comparable sequence read mapping to the frozen control samples.

## Introduction

Parasitic nematodes are important pathogens of humans and other animals. The intestinal soil-transmitted helminths (STHs) – *Ascaris lumbricoides*, hookworms (*Necator americanus* and *Ancylostoma duodenale*), *Trichuris trichiura* and *Strongyloides stercoralis* – infect more than a billion people, with infection concentrated in the young and poor of the tropical developing world (Savioli & Albonico, 2004; Jourdan *et al*., 2018). STH infections are classified by the World Health Organization (WHO) as a Neglected Tropical Disease, because of their substantial burden of infection and public health impact (WHO 2026). Because of the widespread harm that STH infections cause, mass drug administration (MDA) programmes have been implemented in many endemic areas to reduce the prevalence and intensity of STH infection (Bundy *et al*., 2018). Despite these MDA programmes, STHs remain prevalent, and there is the growing concern that they are exerting a strong selective pressure on the STHs, affecting genetic diversity, and potentially leading to the evolution of anthelminthic resistance.

In recent years, great advances have been made in understanding the genomics of the STHs and other parasitic nematodes (Doyle, 2022). Genome assemblies are now available for many of these species and these are now being exploited in a number of ways; for example, to understand their population genetics, pattens of transmission, parasite species identity and host range, and anthelminthic resistance. A range of analyses are used, including PCR-based analyses of single, or relatively few, loci and whole genome sequencing (WGS). Because, by definition, the adult stages of parasitic nematodes live inside their hosts, their transmission stages – commonly eggs or larvae in faeces – are often used as the source of DNA for subsequent molecular analyses. Therefore, how these stages are stored is an important consideration for preserving DNA integrity for down-stream molecular analyses. Successful WGS requires a sufficient quantity and quality of DNA, the latter typically assessed by the absence of contaminating molecules, and the size distribution of DNA molecules. Comparison of different storage conditions for a range of vertebrate tissues showed that low temperature storage (with flash freezing being optimal) was the strongest predictor of the amount of high molecular mass DNA (likely suitable for WGS) that was subsequently extracted (Dahn *et al*., 2022).

Sampling parasitic nematode transmission stages from infected people often requires working in settings where freezing samples is difficult, if not impossible. Therefore, there would be many advantages to ambient temperature storage methods that preserve the samples’ DNA for successful, subsequent analyses.

Ambient temperature storage options include ethanol, Dimethyl sulphoxide-EDTA-saturated sodium chloride (DESS) buffer, and FTA (Flinders Technology Associates) cards. A comparison of ambient temperature ethanol or DESS for storage of a range of eukaryotes (including nematodes) showed that DESS successfully preserved high molecular mass DNA (Ogiso-Tanaka *et al*., 2025). Canine hookworm eggs have also been successfully stored (as judged by PCR amplification of two loci) in ambient temperature DESS (Chen *et al*., 2025). Storage of *S. stercoralis* larvae in ambient temperature ethanol resulted in DNA that could be used for PCR and WGS, though the latter was of variable quality, as judged by the proportion of sequence reads that mapped to the reference genome (Zhou *et al*., 2019).

FTA card storage of blood containing microfilariae has successfully been used for PCR-based detection of filarial nematodes (Duscher *et al*., 2009; Silbermayr *et al*., 2014; Medeiros *et al*., 2015). A study that stored individual eggs or larvae of seven different nematode species on FTA cards, from which DNA was extracted and then used for WGS, found that this was suitable for WGS (measured as the proportion of reads that mapped to the relevant, reference genome) for many, but not all, species and that this variability in-part depended on the life cycle stage being used (Doyle *et al*., 2019). Relevant to the present study, there were fewer mapped sequence reads from infective larvae of *Ancylostoma caninum* and *S. stercoralis*, compared with eggs or other life cycle stages of these species (Doyle *et al*., 2019). Nematode cuticles are collagen-rich structures, with different mixtures of collagen molecules in each life cycle stage. It is possible that the cuticle of infective larvae of parasitic nematodes is particularly strong and resistant to disruption, which may hinder DNA extraction from these stages (Dawkins & Spencer, 1989). FTA card storage has also been used for plant parasitic nematodes, where extracted DNA was suitable for PCR (Marek *et al*., 2014; Peng *et al*., 2017; Camacho *et al*., 2024).

Beyond sample storage, the method of DNA extraction can also affect the quantity and quality of DNA that is obtained. For example, comparison of DNA extraction methods from museum-stored (in ethanol at -20°C for at least 5 years) animal material (though not nematode) showed that a range of DNA extraction methods resulted in DNA that could be used in PCR reactions, though a Cetyltrimethylammonium bromide (CTAB) phenol-chloroform extraction method was optimal (Schiebelhut, *et al*., 2017). Several studies have focused on DNA extraction for parasitic nematodes. A comparison of four commercially-available DNA extraction kits found variable efficacy (as measured by DNA quantification and suitability for PCR analysis) in extracting DNA from Anisakidae nematodes (Seesao *et al*., 2014). For *Teladorsagia circumcincta*, a comparison of 11 DNA extraction methods also showed variable efficacy (again measured by DNA quantification and suitability for PCR analysis) (Sloan *et al*., 2021); similar variability was found when comparing three kits for use with strongyle eggs (Högberg *et al*., 2022).

In preparation for a WGS-based study of hookworms infecting people in Southeast Asia, we wanted to investigate whether there was a suitable ambient temperature storage method for nematode infective third stage (iL3s) that could be used for settings where it would not possible to immediately freeze hookworm larvae. To do this, we compared two ambient temperature storage methods – FTA cards and DESS buffer – for iL3s of two rat-infecting species of nematode, *Nippostrongylus brasiliensis* and *Strongyloides ratti*, seeking to mimic human-infecting nematode samples.

## Materials and Methods

### Nematodes

We used iL3s of the parasitic nematodes of rats, *N. brasiliensis* and *S. ratti*. For both, faecal cultures were made from rats separately infected with each nematode species and the iL3s were recovered in water. Rats were infected with *N. brasiliensis* and *S. ratti* under the authority of licences issued by the Animals (Scientific Procedures) Act 1986.

### Experimental design

We did two experiments where we compared an ambient temperature storage method with frozen storage as the control. Specifically, Experiment 1: iL3s stored at ambient temperature on FTA cards, with frozen iL3s; Experiment 2: iL3s stored in DESS buffer at ambient temperature, with frozen iL3s. In both experiments, the frozen iL3s were the control for the experiment (**Table 1**). In both experiments, for each treatment (FTA / DESS and frozen) there were ten single iL3s, two groups of ten iL3s, and two groups of 50 iL3s.

**Table 1.**
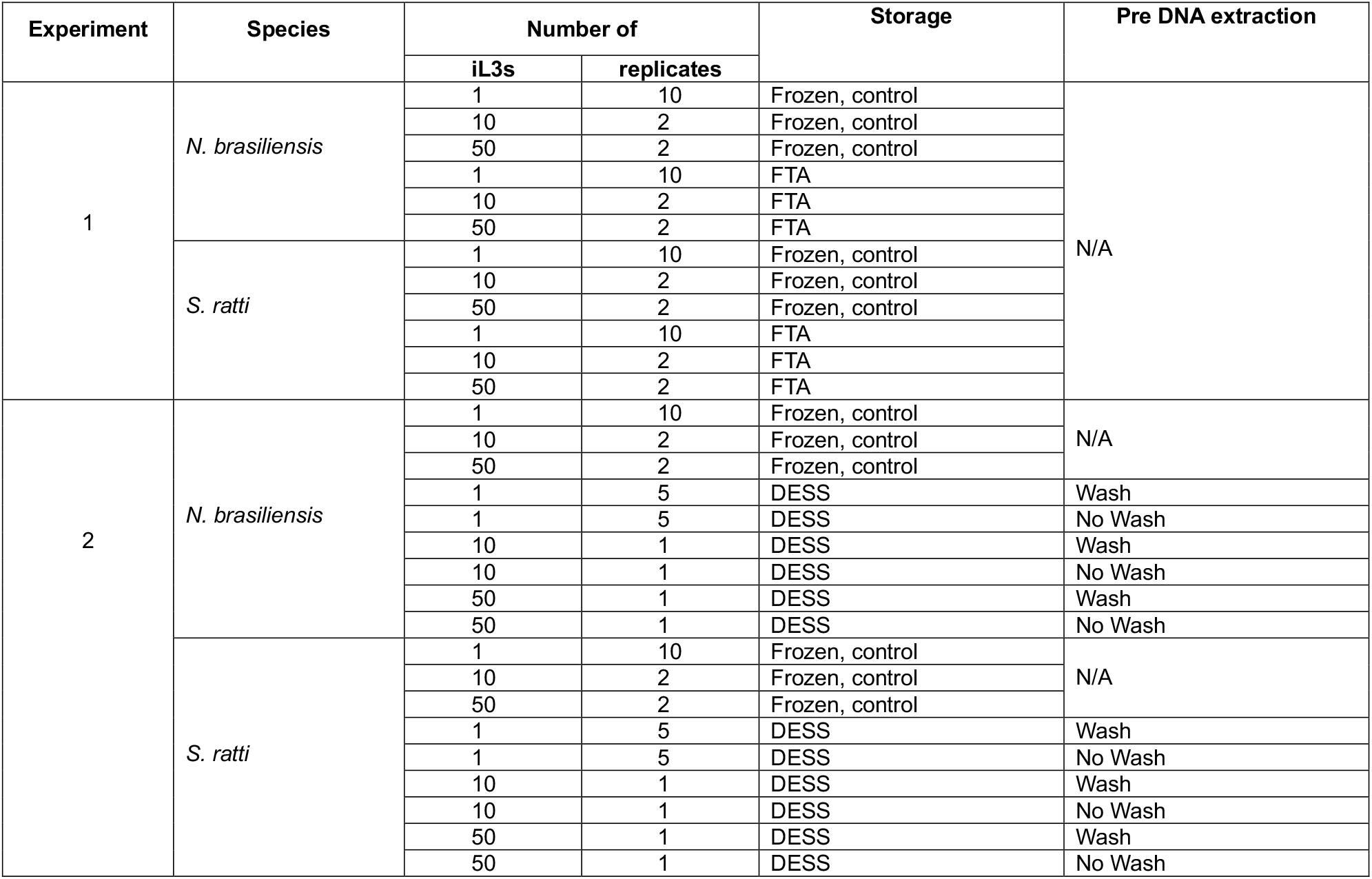
The design of 2 experiments comparing iL3s stored on FTA cards or in DESS buffer with frozen control iL3s. The Pre DNA extraction shows for the DESS-stored samples the Wash and No Wash treatments before DNA extraction.

### Experiment 1: Storage on FTA cards

For the FTA card experiment, single iL3s, groups of ten iL3s, or groups of 50 iL3s were spotted (with a micropipette) separately onto FTA cards (QIAcard FTA CloneSaver, Qiagen) and maintained at ambient temperature for at least 5 days. For the frozen iL3 control, single iL3s, groups of ten iL3s, or groups of 50 iL3s were placed in separate Eppendorf tubes in 10 μL (for single or groups of ten iL3s) or 50 μL (for groups of 50 iL3s) of water, respectively, which were then immediately frozen at -80°C.

To prepare DNA from the FTA card-stored samples, we punched 6 mm diameter holes from the centre of each spotted sample into Eppendorf tubes to which we added 50 μL of 30 mM Tris-HCl (pH 8.0), 0.5 % v/v Tween 20, 0.5 % v/v NP40, 1.25 ug/mL Proteinase K, and then maintained the tubes at 50°C for 1 hour with gentle agitation, and then at 75°C for 30 minutes, centrifuged the tubes and removed the supernatant to fresh tubes, which was then stored at -20°C. To prepare DNA from the frozen control samples we followed a previously described method (Liu *et al*., 2025). Briefly, to each tube we used water to adjust the volume to 50 μL after which we added 70 μL lysis buffer (340 mM NaCl, 170 mM Tris-HCl (pH 8.5), 85 mM EDTA (pH 8), 0.85 % w/v SDS, 1.53 mg/mL Proteinase K, 76.5 mM dithiothreitol), maintained the tubes at 60°C for 2 hours, and then at 85°C for 15 minutes. The tubes were then stored at -20°C.

### Experiment 2: Storage in DESS

For the DESS buffer experiment, single iL3s, groups of ten iL3s, or groups of 50 iL3s were transferred to 100 μL DESS buffer and maintained at room temperature for at least 13 days, after which they were stored at -20°C until processing. DESS buffer was prepared as described by Chen *et al*., 2025. The frozen iL3 controls were prepared exactly as described for Experiment 1 (above).

To prepare DNA from the DESS-stored samples, for half of the samples we reduced the volume of DESS to approximately 10 μL, and then added water to a final volume of 100 μL, this is the “Wash” treatment; the other half of the samples were left in the DESS, which is the “No Wash” treatment (**Table 1; Supplementary Table 1**). For the control samples, water was added to a total volume of 100 μL. For the DESS “Wash”, DESS “No Wash” and control tubes we then added 100 μL lysis buffer to achieve a final concentration of 200 mM NaCl, 100 mM Tris-HCl, 50 mM EDTA, 0.5% w/v SDS, 0.9 mg/mL Proteinase K, 45 mM dithiothreitol. The tubes were then briefly vortexed, centrifuged, and then maintained them at 60°C for 2 hours, and then at 85°C for 15 minutes. To extract DNA from the lysate an Ampure XP DNA cleanup was performed, with a final elution in 40 μL of water.

### Sequencing

For Experiment 1, DNA was purified using Mag-Bind TotalPure NGS Beads (Omega Bio-Tek) and libraries were prepared using the NEBNext Ultra II FS DNA Library Prep Kit for Illumina (New England Biolabs) at half-volume reactions. PCR enrichment was carried out using unique dual indexes (Integrated DNA Technologies) with 13 amplification cycles, purified using Mag-Bind beads and pooled in equimolar amounts and sequenced on the Illumina NovaSeq X Plus platform (Illumina, San Diego, USA) on one lane of a 25B flow cell generating 2 × 150 bp paired-end reads. For Experiment 2, dual-indexed Illumina sequencing libraries were prepared, pooled into a single 56-sample multiplex, and sequenced using 150 bp paired-end reads on a single lane of an Illumina NovaSeq 6000. The sequence data for both experiments are available at the ENA BioProject ERP187659.

### Bioinformatic analyses

Sequencing reads were mapped to the relevant reference genome using a Nextflow mapping pipeline (mapping-helminth/v1.1.2). The reference genomes for both *N. brasiliensis* (PRJNA994163) and *S. ratti* (PRJEB125) were retrieved from WormBaseParaSite V19 (Howe *et al*., 2017). In brief, raw reads were converted into an unmapped BAM (uBAM) file using FastqToSam and adapters were marked using GATK v.4.1.4.1 MarkIlluminaAdapters (McKenna *et al*., 2010). Reads were then mapped to either the *N. brasiliensis* or *S. ratti* reference genome using minimap2 v.2.16 (Li, 2018), after which the mapped reads were sorted using samtools and duplicate reads marked using sambamba (Tarasov *et al*., 2015). Mapping statistics were generated using samtools flagstats and summarised using MultiQC (Ewels *et al*., 2016). The code used to analyse the raw data and to generate the figures is available from GitHub https://github.com/WORMS-SEA-Consortium/FTA_DESS_storage_paper.

## Results

### FTA cards

Storage of individual *N. brasiliensis* or *S. ratti* iL3s on FTA cards resulted in a low proportion of sequence reads successfully mapping to the reference genome (16.6 ± 9.9, 5.6 ± 2.9 %, mean ± SE for *N. brasiliensis* and *S. ratti*, respectively), compared with the frozen control larvae (80.8 ± 0.79, 23.5 ± 1.4 %, mean ± SE for *N. brasiliensis* and *S. ratti*, respectively) (**Figure 1; Supplementary Table 1**). For individual iL3s on FTA cards, only *c*.50-60 % of the number of sequence reads were obtained, compared with the control frozen larvae (101 *vs*. 163 and 77 *vs*. 153 million reads for FTA *vs*. control for *N. brasiliensis* and *S. ratti* respectively; **Supplementary Table 1**). With larger numbers of iL3s, the proportion of reads that mapped to the reference genome improved and was generally comparable to that in the frozen controls. Control, frozen *N. brasiliensis* and *S. ratti* individual iL3s differed, with a much higher proportion of reads mapping for *N. brasiliensis* than for *S. ratti*.

**Figure 1.**
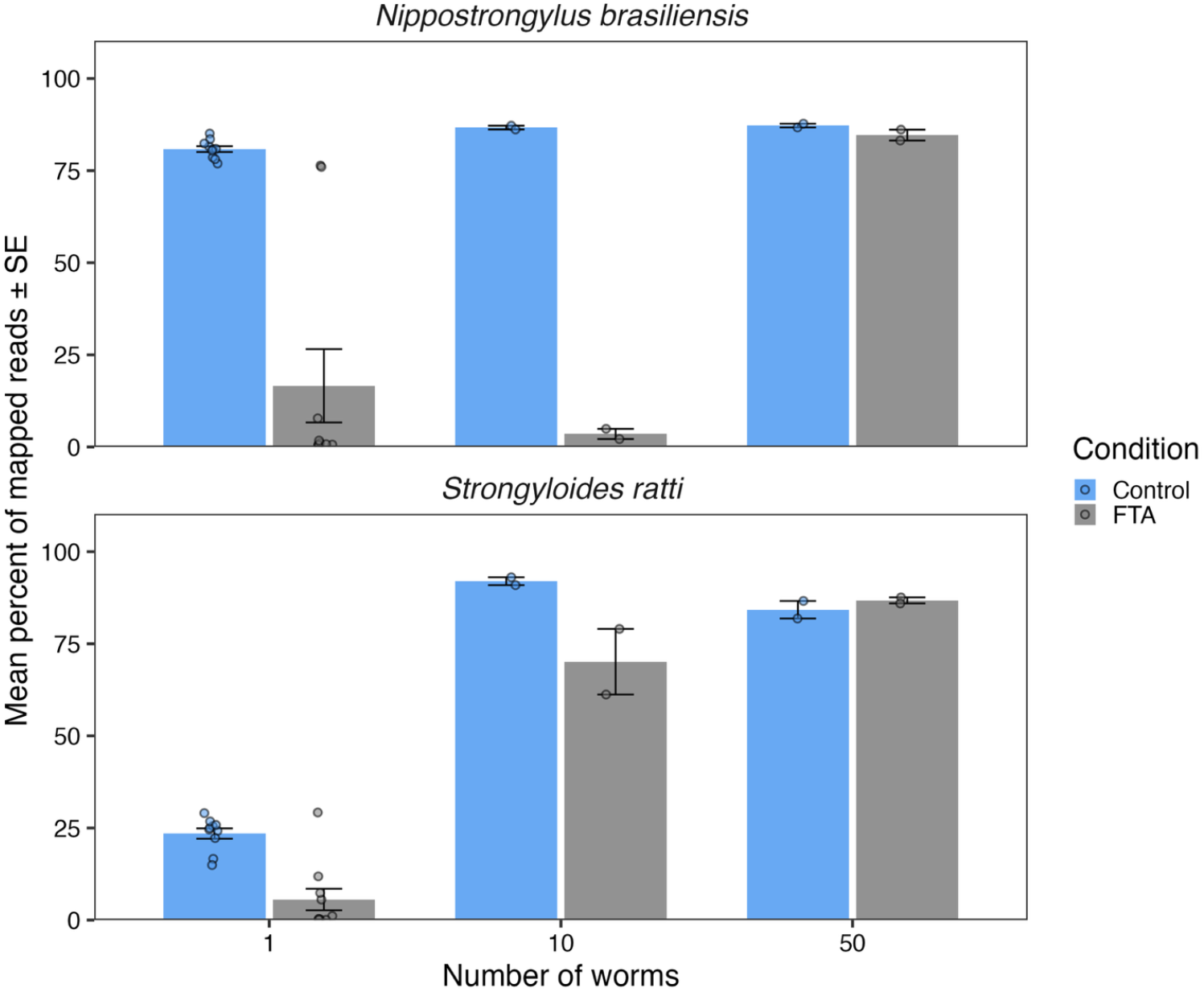
FTA storage. The mean percent of sequence reads that mapped to the *N. brasiliensis* and *S. ratti* genome comparing FTA-storage to frozen control larvae. Error bars are ± 1 SE; individual data points are shown overlaying the bars.

### DESS buffer

Storage of individual *N. brasiliensis* iL3s in DESS (both Wash and No Wash treatments), resulted in a lower and more variable proportion of sequence reads mapping to the reference genome, compared with the frozen control larvae (25.4 ± 9.9, 35.6 ± 6.16, 43.1 ± 1.5 %, mean ± SE for Wash, No Wash and control larvae, respectively) (**Figure 2; Supplementary Table 1**). A comparable number of sequence reads were obtained for individual *N. brasiliensis* iL3s stored in DESS or frozen (**Supplementary Table 1**).

**Figure 2.**
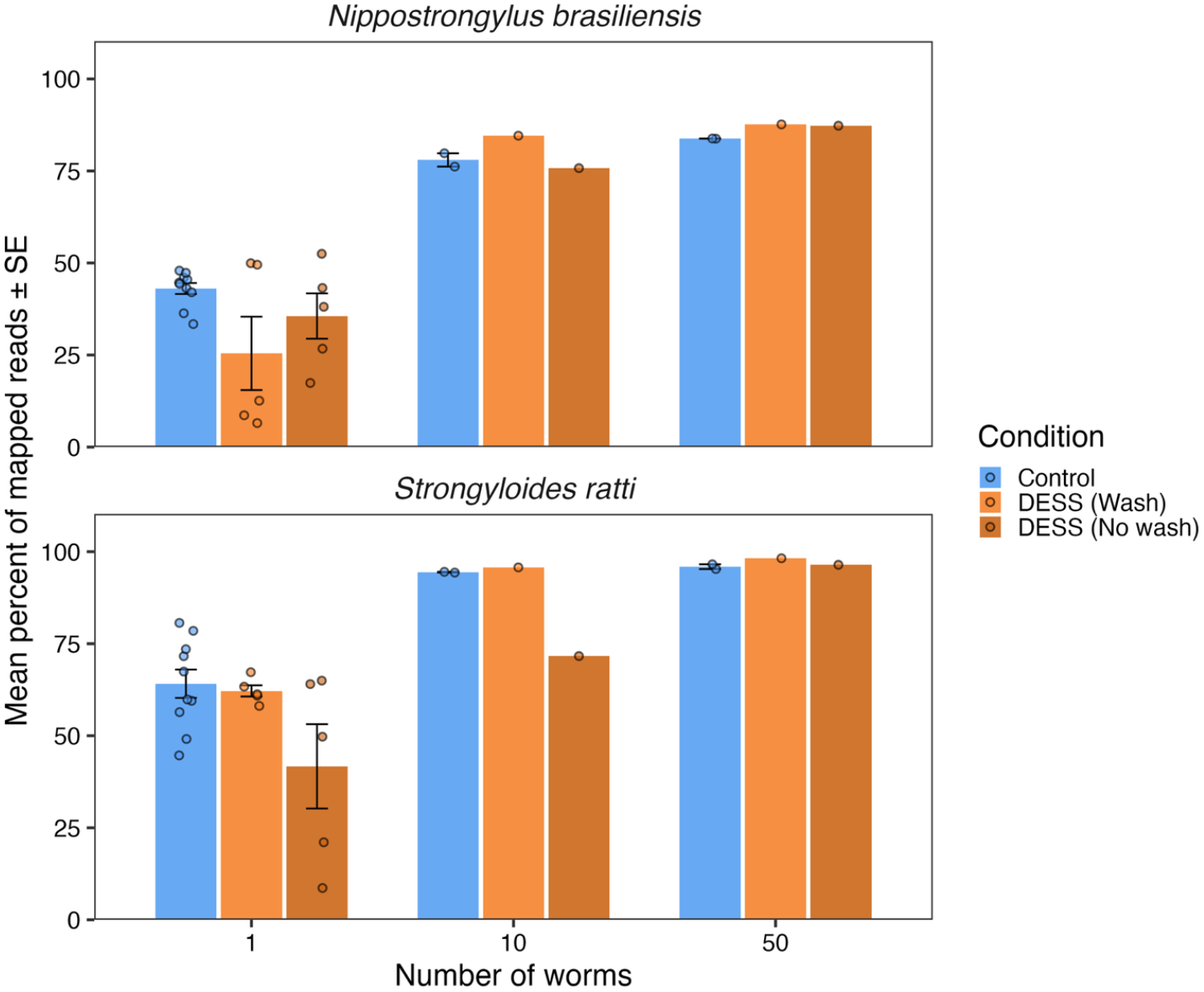
DESS storage. The mean percent of sequence reads that mapped to the *N. brasiliensis* and *S. ratti* genome comparing DESS-storage to frozen control larvae. Error bars are ± 1 SE individual data points are shown overlaying the bars. For the Wash and No Wash treatment for the 10 and 50 larvae samples, there is only a single sample and so no error bar is shown.

Storage of individual *S. ratti* iL3s in DESS (with the Wash treatment before DNA preparation) resulted in the same proportion of sequence reads successfully mapping as the frozen control larvae (**Figure 2; Supplementary Table 1**). There was a slightly lower proportion of reads mapping in the No Wash treatment (62.2 ± 1.5, 41.7 ± 11.5, 64.1 ± 3.8 %, mean ± SE, for Wash, No Wash and control larvae, respectively). A comparable number of sequence reads were obtained for individual *S. ratti* iL3s stored in DESS or frozen (**Supplementary Table 1**).

For groups of 10 or 50 *N. brasiliensis* and *S. ratti* iL3s, the proportion of sequence reads that successfully mapped to the reference genome was comparable between the DESS storage (both with the Wash and No Wash treatment) and the frozen control.

## Discussion

The aim of this study was to test the utility of ambient temperature storage of individual parasitic nematode iL3s for the subsequent recovery of DNA for WGS. The results showed that for individual iL3s, storage on FTA cards or in DESS performed poorly compared with the frozen controls, though DESS storage was better than FTA card storage. However, for groups of 10 or 50 iL3s both ambient temperature storage methods generally performed equivalently to the frozen controls. The exception to this was groups of 10 *N. brasiliensis* iL3s, where the FTA card storage resulted in comparatively fewer sequence reads mapping to the reference genome. Based on these results we conclude that these two ambient temperature storage methods are sub-optimal for routine, field-based storage of single nematode iL3s for subsequent WGS, especially if immediate frozen storage is available. However, these ambient temperature storage methods are suitable for larger groups of iL3s when more DNA is available.

The ambient temperature stored larvae resulted in a lower proportion of reads mapping to the reference genomes compared with the frozen control larvae. This is consistent with ambient temperature storage of individual larvae resulting in comparatively more non-nematode DNA, and so lower mapping rates. FTA storage also resulted in fewer sequence reads overall, compared with the frozen control larvae. This is consistent with comparatively lower quantities of DNA and / or poor quality DNA resulting from the FTA storage. The relative success of both ambient temperature storage methods for groups of 10 or 50 iL3s is consistent with there being less non-nematode DNA contamination, resulting in higher mapping rates.

The higher mapping rates of the 10 and 50 iL3 samples, compared to that of the individual iL3s, suggests that more starting DNA improved the likelihood of sequencing success. For individual iL3 samples, whole genome amplification (WGA) methods could be used with ambient temperature stored individual iL3s, though the utility of these methods also depends on the quantity and quality of input DNA (Hshieh & Maloney, 2025). While our results show that ambient temperature storage of groups of 10 or 50 iL3s is suitable for WGS, whether such pooled samples are useful depends on the scientific questions that are being asked. For example, for population genetic studies, analysis of individual samples is often ideal since this can allow a range of analyses of individual genotypes. Analyses of pooled samples can be more limited, and is often based on the inference of allele frequencies. However, sequencing of pooled samples has been used in single nucleotide polymorphism (SNP) discovery and in genome-wide scans of selection (Babineau *et al*., 2022; Doyle *et al*., 2022).

As we review in our Introduction, FTA cards and DESS buffer have previously been used to store nematode samples, though the efficacy of this storage was mainly judged by PCR analysis, rather than WGS. In the studies where WGS was used the results were, (i) variable (Zhou *et al*., 2019), and (ii) appeared to depend on the life cycle stage being used (Doyle *et al*., 2019). Our results also show this inter-individual variability in the proportion of successfully mapped reads. Regarding the life cycle stage, we used iL3s, which is also consistent with the reported comparatively fewer mapped reads of infective larvae of *A. caninum* and *S. stercoralis* (Doyle *et al*., 2019).

*N. brasiliensis* and *S. ratti* have different genome sizes, *c*.290 and *c*.43 Mb, respectively, and different GC richness *c*.42% and *c*.21%, respectively (WormBaseParaSite.org; Hunt *et al*., 2016). However, these different genome characteristics are unlikely to explain the differences in the efficacy of FTA or DESS storage between the two species. Human-infecting hookworms and *N. brasiliensis* are phylogenetically more closely related to each other than to *S. ratti*, and genomes of the human-infecting hookworms are comparatively more similar to that of *N. brasiliensis*; specifically, 244 and 332 Mb, 40 and 42% GC for *N. americanus* and *A. duodenale*, respectively (Tang *et al*., 2014; WormBaseParaSite.org) suggesting that *N. brasiliensis* is the more appropriate model for DNA and genomic studies of human hookworms.

Our study has a number of limitations. First, with our focus on ambient temperature storage of individual iL3s, we used a relatively small number of replicates in the Experiment 2 Wash and No Wash treatments. It would be beneficial in a future study to use more replicates to better measure the efficacy of the Wash and No Wash treatments on the proportion of successfully mapped sequence reads. Second, the rationale of our study was to test methods for eventual use with human-infecting hookworms. However, we used iL3s of two species of rat parasites, and it is possible that different results would be obtained with *N. americanus* and *A. duodenale*. Third, we only tested the ambient temperature storage of iL3s, and other life cycle stages may store differently on FTA cards or in DESS buffer. Fourth, we did not test different lengths of ambient temperature storage, and it is possible that with shorter storage better WGS results would have been obtained. Fifth, the frozen iL3 controls of the two experiments differed in the proportion of sequence reads that mapped to the relevant, reference genome. Specifically, for *S. ratti* single, frozen control iL3s, the proportion of successfully mapped reads was 23.5 and 64.3 % in Experiments 1 and 2, respectively; for *N. brasiliensis* this was 80.8 and 43 % in Experiments 1 and 2, respectively. Because of this, the results of the two experiments cannot be directly compared.

In conclusion, ambient temperature storage of individual iL3s of *S. ratti* and *N. brasiliensis* on FTA cards or in DESS buffer is less effective than frozen storage for subsequent WGS. However, given that the DESS ambient temperature storage of individual iL3s did result in substantial proportions of successfully mapped sequence reads, we suggest that in settings where immediate frozen storage is not possible, then DESS storage could be used as a temporary alternative.

## Acknowledgments

We would like to thank Tom Bosley, Jordan Boucher and Brian Chan for help in providing nematode larvae This work was funded by MRC grant UKRI989, the University of Liverpool, the Wellcome Trust (UK) through core funding to the Wellcome Sanger Institute (UK) (220540/Z/20/A), and a UKRI Future Leaders Fellowship (MR/T020733/1).

The WORMS-SEA consortium is: in Indonesia Erwan Budi Hartadi and Elsa Herdiana Murhandarwati; in Malaysia, Hesham Mahyoub Al-Mekhlafi; and Arutchelvan Rajamankam; in the Philippines June Haidee Acūna-Lariosa, Jennifer Luchavez and Vanessa Joy Mapalo; and in the UK Modupeh Betts, Stephen Doyle and Mark Viney.

## Author contribution, by CRediT taxonomy

**Erwan Budi Hartadi:** Conceptualization, Funding acquisition, Writing – review and editing

**Elsa Herdiana:** Conceptualization, Funding acquisition, Writing – review and editing

**Hesham Mahyoub Al-Mekhlafi:** Conceptualization, Funding acquisition, Writing – review and editing

**Arutchelvan Rajamankam:** Conceptualization, Funding acquisition, Writing – review and editing

**June Haidee Acūna-Lariosa:** Conceptualization, Funding acquisition, Writing – review and editing

**Jennifer Luchavez:** Conceptualization, Funding acquisition, Writing – review and editing

**Vanessa Joy Mapalo:** Writing – review and editing

**Modupeh Betts:** Writing – review and editing

**Stephen Doyle:** Conceptualization, Data curation, Formal Analysis, Funding acquisition, Investigation, Writing – review and editing

**Mark Viney:** Conceptualization, Funding acquisition, Investigation, Writing – original draft preparation

